# A role for aquaporin (Aqp1) in the control of *Cryptococcus neoformans* cell morphology

**DOI:** 10.64898/2026.01.29.702379

**Authors:** Piotr R. Stempinski, J. Alberto Patiño-Medina, Francisco G. Hernandez, Isabel A. Jimenez, Samuel Rodrigues dos Santos Junior, Peter Agre, Arturo Casadevall

## Abstract

Aquaporins are small, integral membrane channels that facilitate the transport of water across cellular membranes and, in the case of aquaglyceroporins, can also conduct specific neutral solutes, such as glycerol. These proteins are conserved across biological kingdoms, yet their roles in fungal virulence remain relatively understudied. In *Cryptococcus neoformans*, an opportunistic fungal pathogen, we examined the organism’s single aquaporin, Aqp1, and uncovered unanticipated influences on cellular morphology. Loss of Aqp1 resulted in smaller cells, whereas its presence promoted the formation of enlarged titan-like cells. This shift in size was closely linked to intracellular redox physiology. Consequently, the overexpression of the cryptococcal aquaporin increased sensitivity to oxidative stress and led to the largest titan-like cells; antioxidant supplementation suppressed this enlargement, consistent with a ROS-dependent regulatory mechanism. Additionally, Aqp1 overexpression produced vacuolar abnormalities in titan-like cells, suggesting that excessive water influx strained intracellular organization during rapid cell expansion. These findings position Aqp1 at a functional crossroads connecting membrane transport, oxidative balance, and size control, and they support a model in which an aquaporin contributes to the morphological plasticity that allows *C. neoformans* to adapt to environmental pressures.

## Introduction

Aquaporins are a conserved family of transmembrane protein channels that govern the transport of water and, in some cases, small molecules such as glycerol, urea, and ammonia across biological membranes (1–4). These channels play important roles in maintaining osmotic balance, regulating intracellular turgor pressure, and mitigating response to external stress factors (1, 2, 5, 6). Aquaporins can be categorized into two classes: orthodox aquaporins, which are highly selective for water, and aquaglyceroporins, which allow the transport of water and some small uncharged molecules (1, 5, 7, 8). This functional diversity reflects their wide-ranging physiological roles across all kingdoms of life (3, 6, 7, 9–11). In fungi, aquaporins have been implicated in development, morphogenesis, freeze tolerance, sporulation, and responses to osmotic and oxidative stress (12–14). However, the roles of aquaporins in regulating virulence-associated processes of pathogenic fungi remain largely unexplored.

*Cryptococcus neoformans* is an opportunistic fungal pathogen and the etiologic agent of cryptococcosis, which is a life-threatening pulmonary and central nervous system infection, particularly in immunocompromised individuals (15–19). Cryptococcosis accounts for an estimated 180,000 deaths annually, disproportionately affecting populations with high prevalence of advanced HIV infection. The success of *C. neoformans* as a pathogen is primarily attributed to its ability to withstand and adapt to hostile host environments through diverse stress response mechanisms, production of multiple virulence factors, immune evasion strategies, and morphological plasticity (16, 20–26).

A key morphological feature of *Cryptococcus* is a polysaccharide capsule, a major virulence factor that surrounds both typical and morphologically specialized yeast cell forms (19, 27–29). The capsule plays a critical role in protecting the fungus from host immune defenses, facilitating immune evasion and persistence during infection (19, 30). Among the various morphological adaptations observed in host cells during infection, the formation of titan cells represents a striking example of cryptococcal plasticity (17, 31). These enlarged, polyploid cells are characterized by thickened cell walls, expanded capsules, and altered surface properties that enhance resistance to phagocytosis and increase tolerance to oxidative and nitrosative stresses (27, 32, 33). Titan cells are thought to modulate host immune responses and contribute to fungal survival and dissemination (27, 34, 35). Despite their importance in the pathogenesis of cryptococcosis, the molecular mechanisms driving titan cell formation remain incompletely understood.

In this study, we characterize the function of Aqp1, the single aquaporin identified in *C. neoformans,* through a combination of phenotypic, morphological, and structural analyses (12). Our studies reveal that Aqp1 can influence cell size and stimulate the formation of titan-like cells in *C. neoformans* in a ROS-dependent manner. Furthermore, we confirmed that Aqp1 participates in maintaining intracellular redox balance and distribution of reactive oxygen species within the cell. This connection between water channel function, oxidative homeostasis, and morphogenesis suggests that aquaporins contribute to the adaptive flexibility that allows *C. neoformans* cells to remodel their cellular morphology and size to form titan cells in response to environmental stressors.

## Results

### Structural prediction and homology analysis of cryptococcal Aqp1

To explore evolutionary conservation and potential functional similarities, we compared the amino acid sequence of *C. neoformans* Aqp1 with human aquaporins. BLASTp analysis identified the highest similarity between cryptococcal Aqp1 and human AQP8, a mitochondrial aquaporin involved in water and small solute transport (Figure 1A)(36–38). AlphaFold predictive modeling of the secondary structure of cryptococcal Aqp1 revealed a classical aquaporin structure composed of six transmembrane helices and two conserved NPA (Asn-Pro-Ala) motifs, consistent with canonical water channel proteins (Figure 1B)(1, 39). ChimeraX structural alignment revealed high structural homology, both in positional alignment and in surface hydrophobicity and electrostatic charges, between the cryptococcal Aqp1 and its human homologs (Figure 1B).

**Figure 1.**
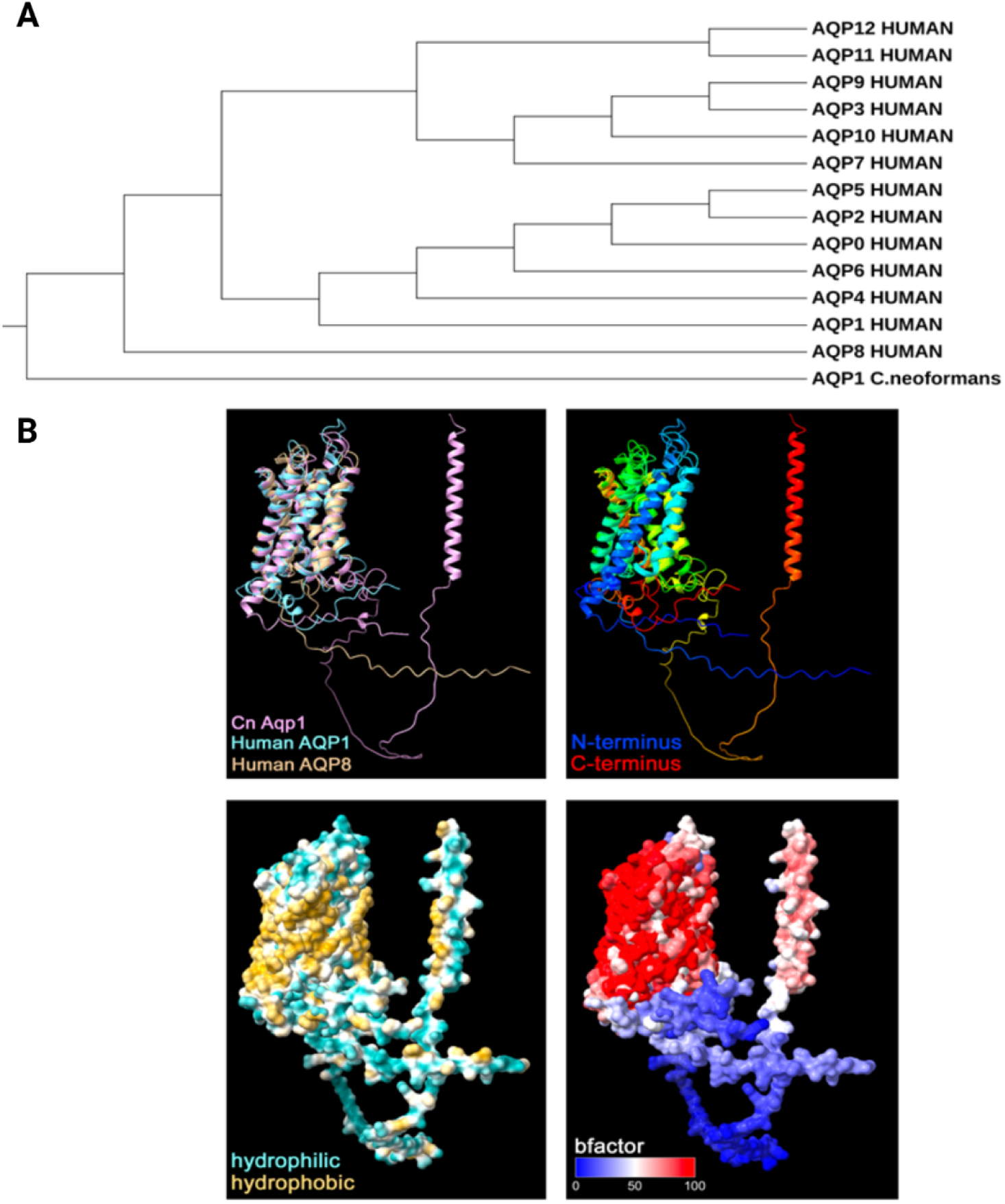
Evolutionary comparison and structural modeling of Cryptococcus neoformans Aqp1. (A) Phylogenetic analysis comparing the amino acid sequence of C. neoformans Aqp1 with human aquaporins, highlighting evolutionary relationships and potential functional similarities. (B) AlphaFold predictive modeling of Aqp1 revealed a classical aquaporin fold, comprising six transmembrane helices and two conserved NPA (Asn-Pro-Ala) motifs, consistent with canonical water channel proteins. Structural alignment using ChimeraX demonstrated high structural homology between cryptococcal Aqp1 and its closest human counterparts, AQP8 and AQP1, including conserved positional alignment and similar surface hydrophobicity and electrostatic profiles.

However, cryptococcal Aqp1 has an additional C-terminal domain that is not present in its human homologs. The existence of the C-terminal transmembrane domain is not conserved, and its presence is variable in different fungal species (Supplementary figure 1). Thus, the predicted structure supports the functional classification of Aqp1 as a member of the aquaporin family, but also reveals an additional domain, absent from human aquaporin homologs, that may have functional significance specific to Cryptococcus spp. Using predictive modeling, this C-terminal domain was predicted to be intracellular, rather than extracellular or transmembrane, and was not predicted to be a signal peptide. To further evaluate the presence of Aquaporin-like domain (PF00230) across clinically relevant fungi, we performed Ortholog group analyses of the proteomes from multiple species. Our analysis revealed considerable variation in the number of Aqp1 homologues among different fungi (Table 1). The presence of a single protein containing an aquaporin domain (PF00230), such as found in *C. neoformans*, is relatively uncommon but not unique, as other species such as *Cryptococcus gattii*, *Malassezia sympodialis*, *Candida parapsilosis*, or *Rhizopus microspores*, also possess only one putative aquaporin. In contrast, several fungal species have more aquaporin homologues, including *Exophiala spinifera* with 13 homologues and *Cladophialophora bantiana* with 10 channels. These findings suggest that aquaporin gene copy number is highly variable across fungal taxa, potentially reflecting differences in ecological adaptation and physiology of diverse fungal species.

**Table 1.** Overview of selected clinically relevant fungal species and Aqp1 homologues. This table lists the clinically relevent fungal species included in our analysis, indicating their taxonomic division, the specific strains examined, and the number of Aqp1 protein homologues identified in each strain.

| Division | Species | Strain | No. of homologues |
| --- | --- | --- | --- |
| **Basidiomycota** | *Cryptococcus gattii* | VGII R265 | 1 |
|  | *Malassezia sympodialis* | ATCC 42132 | 1 |
|  | *Trichosporon asahii* | CBS 2479 | 4 |
| **Ascomycota** | *Aspergillus fumigatus* | Af293 | 3 |
|  | *Aspergillus nidulans* | FGSC A4 | 5 |
|  | *Aspergillus niger* | ATCC 1015 | 7 |
|  | *Blastomyces dermatitidis* | ATCC 26199 | 2 |
|  | *Candida auris* | B11220 | 1 |
|  | *Candida glabrata* | BG2 | 4 |
|  | *Candida parapsilosis* | CDC317 | 1 |
|  | *Cladophialophora bantiana* | CBS 173.52 | 10 |
|  | *Coccidioides immitis* | H538.4 | 2 |
|  | *Emergomyces africanus* | CBS 136260 | 2 |
|  | *Exophiala spinifera* | CBS 89968 | 13 |
|  | *Fonsecaea pedrosoi* | CBS 271.37 | 9 |
|  | *Fusarium solani* | FSSC 5 MPI-SDFR-AT-0091 | 5 |
|  | *Histoplasma capsulatum* | G184AR | 2 |
|  | *Lomentospora prolificans* | JHH-5317 | 4 |
|  | *Microsporum canis* | CBS 113480 | 2 |
|  | *Paracoccidioides brasiliensis* | Pb03 | 2 |
|  | *Sporothrix schenckii* | 1099-18 | 2 |
|  | *Stachybotrys chartarum* | IBT 40293 | 6 |
|  | *Trichophyton rubrum* | CBS 118892 | 2 |
|  | *Verruconis gallopava* | CBS 43764 | 8 |
| **Mucoromycota** | *Rhizopus microsporus* | ATCC 52814 | 1 |
|  | *Mucor circinelloides* | 1006PhL | 2 |

### Growth of *C. neoformans* aquaglyceroporin mutants in media containing glucose or glycerol as the main carbon source

Earlier sequence analysis suggested that Aqp1 in *C. neoformans* could act as an aquaglyceroporin capable of transporting glycerol and other small solutes (12). To confirm this experimentally, the wild-type strain H99, the deletion mutant (*aqp1*Δ), the complemented strain (*aqp1*Δ::AQP1), and the overexpression strain (AQP1 OE) were each tested for growth in media where either glucose or glycerol served as the primary carbon source. In YNB media supplemented with glucose, all tested strains showed similar growth rates over six days of incubation (Figure 2A), implying that Aqp1 was not required for growth when glucose is available. When glycerol replaced glucose in the same type of medium, only the AQP1 OE strain manifested an increased growth rate (Figure 2B). Growth in this condition was delayed and modest compared to glucose-grown cultures, yet the overexpression strain continued to proliferate, unlike H99, *aqp1*Δ, and *aqp1*Δ::*AQP1* strains. Growth differences were also assessed in minimal medium (MM) containing either glucose or glycerol as a primary carbon source. In MM–glucose, all tested strains proliferated at similar rates for eight days (Figure 2C). In the conditions where glycerol was the sole carbon source, a distinct pattern emerged (Figure 2D). Growth was initially slow for all strains, but after about four days, the AQP1 OE strain presented an increase in proliferation rate, reaching the highest cell concentration and showing the steepest increase in growth, followed by H99 and *aqp1*Δ::*AQP1*, which grew at intermediate rates. The *aqp1*Δ mutant proliferated slowly and remained at the lowest cell concentration during the entire incubation period. These findings indicate that Aqp1 facilitates growth when glycerol is the available carbon source. The superior performance of the AQP1 OE strain and the impaired growth of the *aqp1*Δ mutant support the view that Aqp1 functions as an aquaglyceroporin in *C. neoformans*, facilitating glycerol uptake and contributing to metabolic flexibility under nutrient-limited conditions.

**Figure 2.**
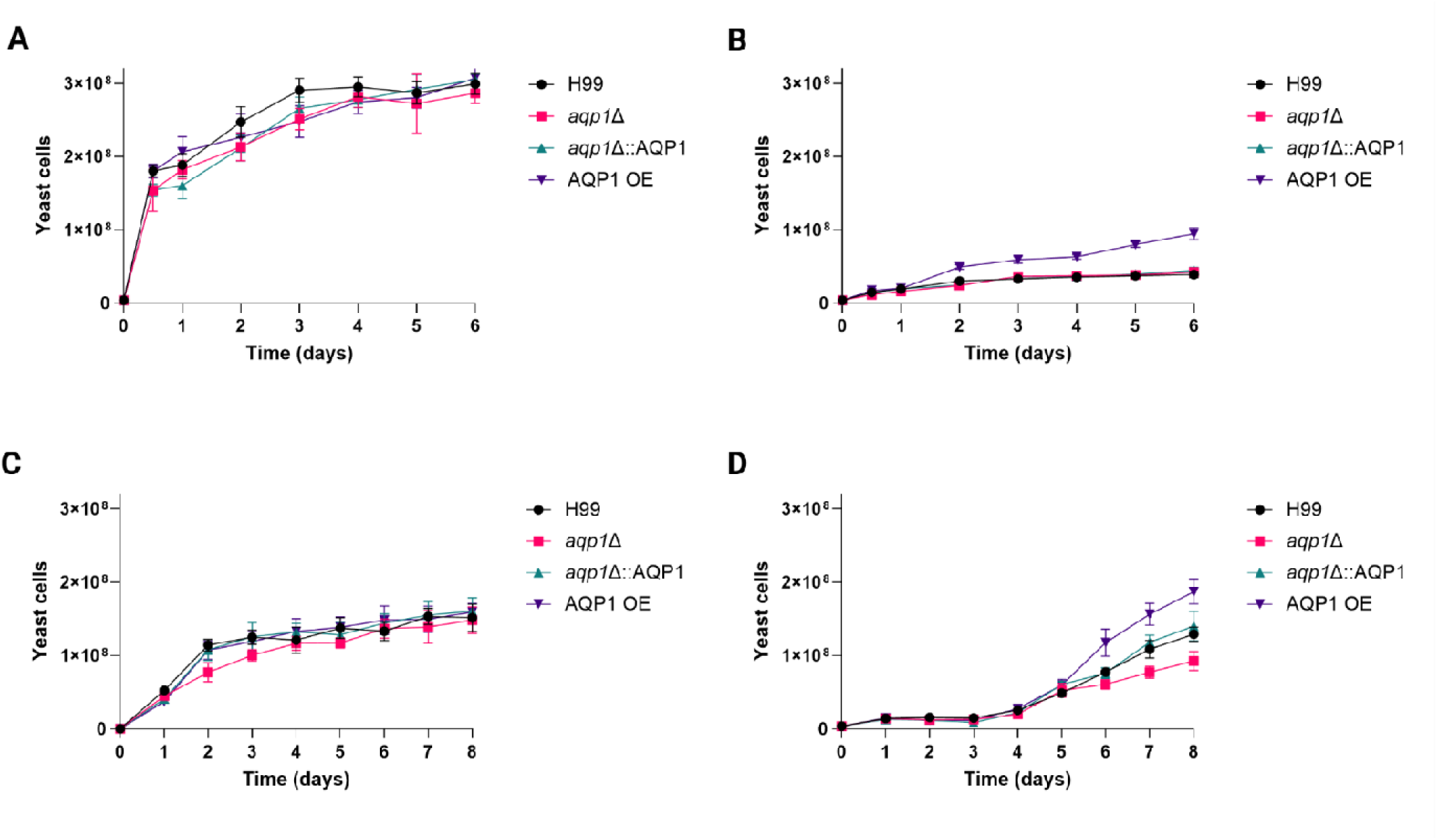
Comparative growth analysis of C. neoformans strains under glycerol as unique carbon source. The figure shows the growth dynamics of four C. neoformans strains: H99, aqp1Δ, aqp1Δ::AQP, and AQP1 OE. Glucose media were used as a control. (A) Growth curves of all strains cultured in yeast nitrogen base (YNB) medium containing 111 mM glucose as the carbon source. Culture were incubated for 6 days, and cell density was measured every 24 hours. (B) Growth curves of all strains grown in YNB medium in which glucose was replaced with 111 mM glycerol as the sole carbon source. (C) Growth curves of the strains incubated in minimal medium (MM) containing 111 mM glucose for 8 days, with cell density measured every 24 hours. (D) Growth curves of the strains grown in MM supplemented with 111 mM glycerol instead of glucose, under identical incubation and measurement conditions. Data represents the mean of three independent biological replicates.

### Aquaporin Aqp1 does not affect the growth of *C. neoformans* under temperature stress, fluconazole treatment or melanization

Aquaporins were previously shown as essential components of high-temperature stress response in various plants, animals, and fungi (9, 40–46). To determine whether Aqp1 contributes to the thermal stress adaptation of *C. neoformans*, we performed spot dilution assays using the wild-type strain H99, an *aqp1* deletion mutant (*aqp1*Δ), a complemented strain (*aqp1*Δ::AQP1), and a strain overexpressing Aqp1 (AQP1 OE).

Ten-fold serial dilutions of each strain were spotted onto YPD agar and incubated at 30°C, 37 °C, or 39 °C to assess growth under normal and elevated temperature conditions. All strains presented similar growth at each temperature (Supplementary figure 2A). To investigate if Aqp1 played a role in low-temperature adaptation, the same set of strains was analyzed on both minimal media (MM) and YPD solid media at 4 °C, and plates were incubated for 6, 12, and 18 days to accommodate slower growth at this temperature (Supplementary figure 2B and 2C). All strains showed similar growth across all tested conditions. This observation suggests that Aqp1 is not required for cellular adaptation to high or low-temperature stress responses in *C. neoformans*. To assess the potential role of aquaporins in antifungal susceptibility, we conducted a dilution spot assay on YPD agar media supplemented with 0, 8, or 12 µg/mL fluconazole. All strains exhibited comparable growth across the tested concentrations (Supplementary figure 2D), confirming that aquaporin function does not influence susceptibility or resistance to fluconazole. Additional evaluation of cryptococcal melanin production was performed by spotting wild type and aquaporin mutant strains onto MM agar plate supplemented with L-DOPA. All tested strains presented a similar rate of melanin production (Supplementary figure 2E).

### Aquaporin Aqp1 does not influence cell size, capsule thickness, or buoyancy in nutrient-rich conditions

To examine the potential role of Aqp1 for cryptococcal morphology or cell density, we analyzed cell size, capsule thickness, and buoyancy in wild-type H99, *aqp1*Δ, *aqp1*Δ::AQP1, and AQP1 OE strains following growth in YPD liquid media for 48 h.

Quantitative measurements revealed no differences in cell size or capsule dimensions among the tested strains (Figure 3A and 3B). To assess potential changes in cell density, we used cell buoyancy as a proxy by measuring the rate of passive cell settling in static liquid cultures, under the assumption that cells with lower density would exhibit more buoyancy. All strains displayed similar buoyancy, with no differences in settling rate between strains (Figure 3C and 3D). Thus, Aqp1 did not significantly alter cell size, capsular size or buoyancy of *C. neoformans* cells under nutrient-rich growth conditions. The assessment of the changes in cell surface hydrophobicity was evaluated using MATH assay, which revealed that the overexpression mutant strain presented a substantial increase in cell surface hydrophobicity (Figure 3E). Additionally, the deletion mutant strain presented a small but significant change in cell surface hydrophobicity, supporting previous suggestions that aquaporin may play a crucial role in processes managing cellular hydrophobicity (12).

**Figure 3.**
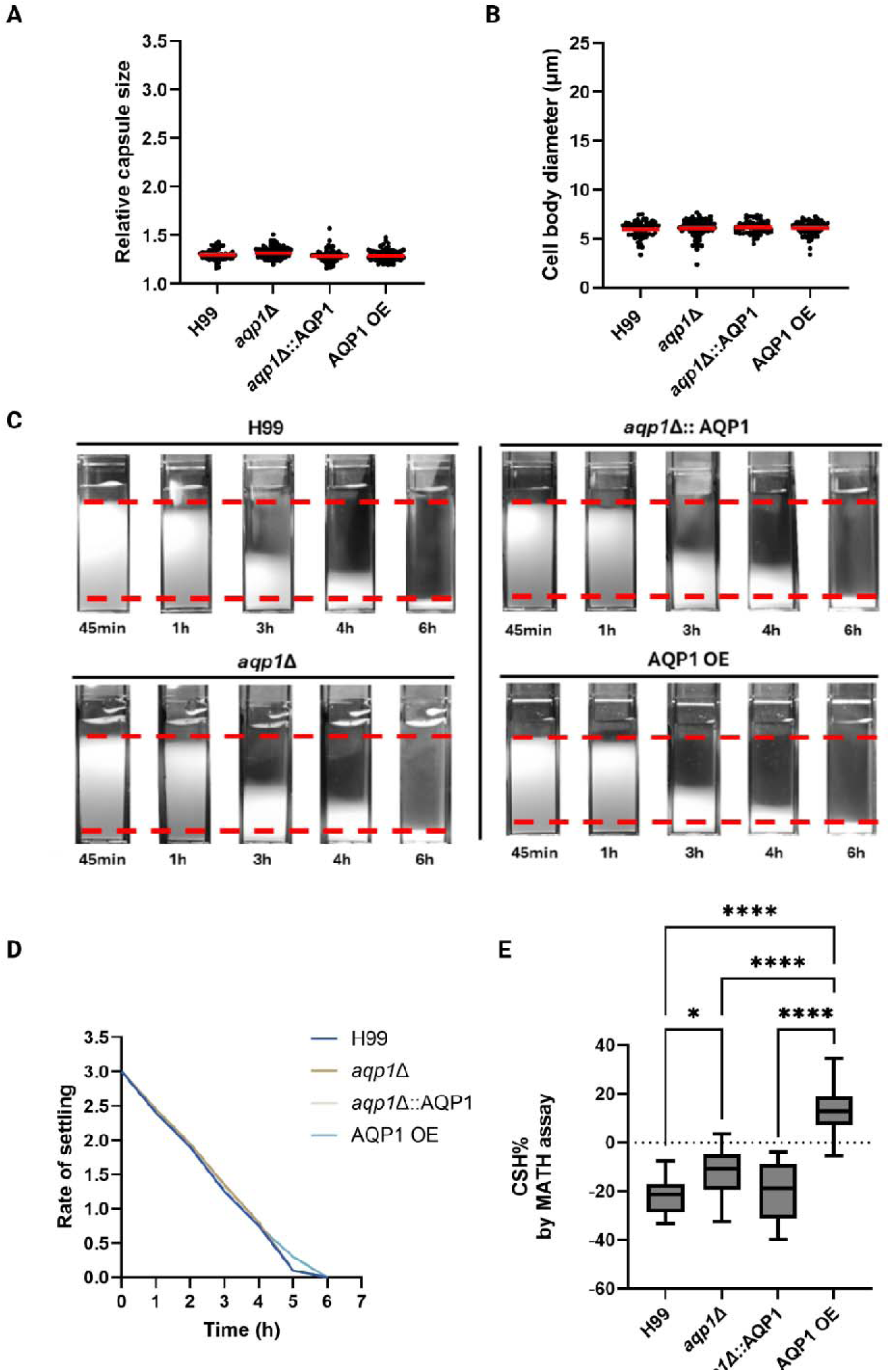
Analysis of cell buoyancy and hydrophobicity in wild-type, aqp1Δ, aqp1Δ::AQP1, and AQP1 OE strains based on capsule size, cell body diameter, and settling speed. (A) Scatter plot showing the relative capsule size for each strain. (B) Scatter plot displaying cell body diameter measurements. Each point represents an individual cell; bars indicate the mean. (C) Representative images from the settling assay for all four strains at various timepoints. Red dotted lines indicate the initial and final positions of the cell suspension during settling. (D) Line graph showing the rate of cell settling over time for each strain. (E) Box plot presenting the level of hydrophobicity measured for each strain. Statistical analysis was performed using 2way ANOVA test.

### Aquaporin overexpression enhances capsule and cell enlargement under capsule-inducing conditions

The cellular morphology of *C. neoformans* is strongly influenced by the surrounding environmental factors and various stress conditions that may modulate its adaptive stress responses (26, 28, 31, 47–49). To evaluate if Aqp1 could modulate cell morphology in capsule-inducing conditions, we cultured wild-type and aquaporin mutant strains in MM at 30°C for 3 days. Under these conditions, the AQP1-overexpressing cells showed increased average cell size and capsule thickness compared to the other tested strains (Figure 4A and 4B). Surprisingly, a substantial subpopulation of cells from the aquaporin overexpression strain exceeded 10 µm in diameter, a phenotype that was not observed in the wild-type or mutant strains (Figure 4C and 4D). Further evaluation of the cell size of tested strains for an extended period (up to 8 days) showed that a small subpopulation of cells in the strain with overexpressed aquaporin 1 exceeding 10 µm (Figure 4E). These findings suggest that elevated Aqp1 expression strongly promotes cellular and capsular enlargement in a small subpopulation of cells exposed to low-nutrient, capsule-inducing conditions.

**Figure 4.**
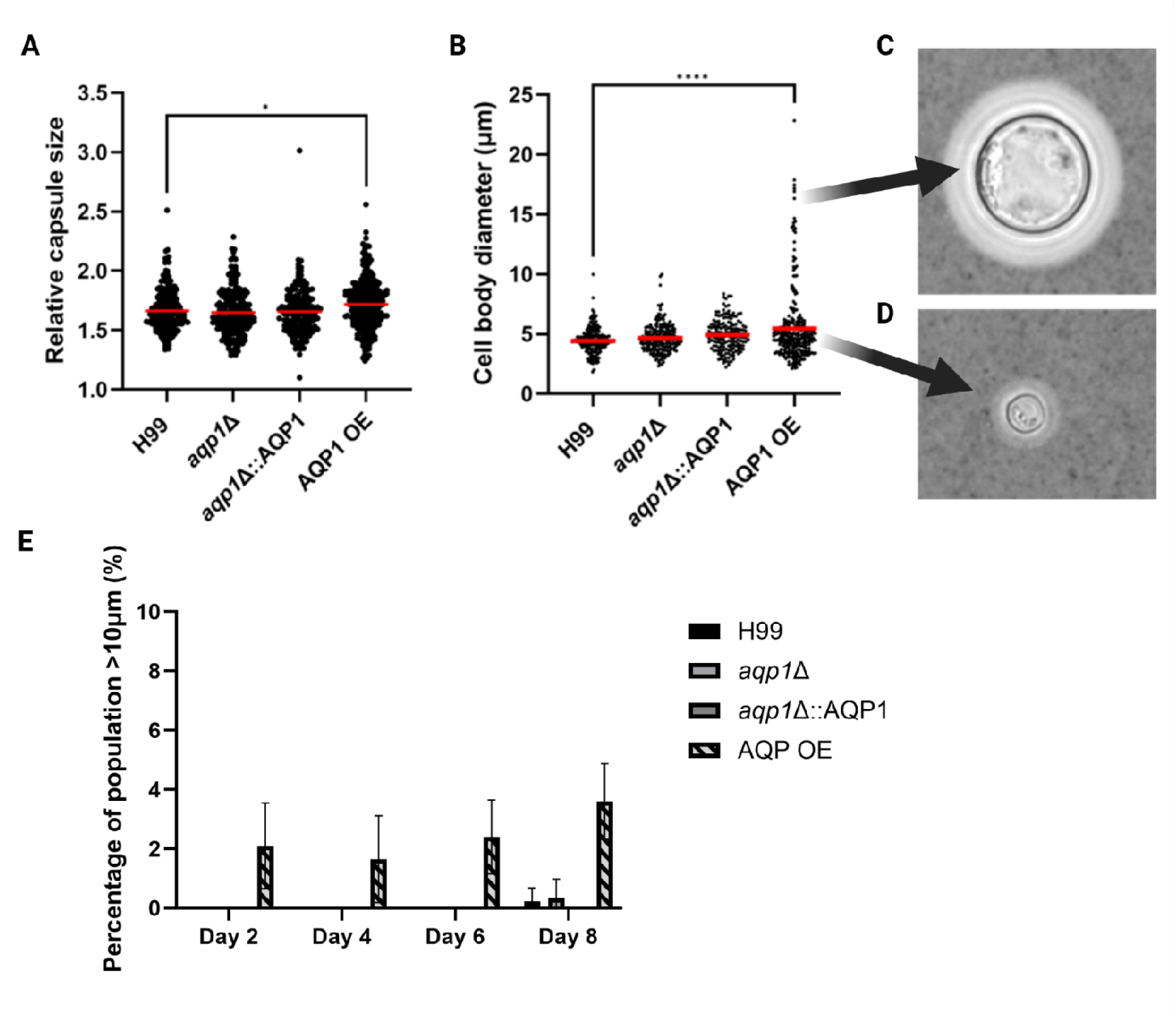
Analysis of polysaccharide capsule size and cell body diameter in wild-type, AQP1 deletion, complemented, and overexpression strains grown in capsule-inducing minimal media. (A) Scatter plot showing relative capsule size for all four strains. (B) Scatter plot displaying cell body diameter measurements across the same strains. Each point represents an individual cell; bars indicate mean ± SD. (C) India ink-stained image of an enlarged cell observed within the AQP1-overexpressing strain population, captured under a 40× objective. (D) India ink-stained image of a representative cell with average size, also imaged under a 40× objective. Panels C and D highlight morphological heterogeneity observed in the AQP1-overexpressing strain. (E) Bar graph representing the mean percentage of cell population exceeding 10 µm in diameter measured after 2, 4, 6, and 8 d of incubation in MM at 30 °C. Data represent the mean from three independent biological replicates. Statistical analysis was performed using 2way ANOVA test.

### Aqp1 influences cell size in low-inoculum cultures grown under capsule-inducing conditions

To further explore the role of Aqp1 in cryptococcal morphogenesis under capsule-inducing conditions, we inoculated wild-type and aquaporin mutant strains into minimal medium at a low starting density (1,000 cells per mL of culture) and incubated them at 30 °C for 4 d. Cell body and capsule sizes were then assessed by India ink staining and light microscopy. The AQP1-overexpressing strain formed cells with modest but statistically significant increases in capsule thickness compared to the wild-type strain (Figure 5A). Additionally, the *aqp1*Δ strain displayed a significantly reduced average cell body diameter (∼5 µm), while WT, complemented, and overexpression strains showed a marked increase in cell size (Figure 5B). In these conditions, the aquaporin deletion mutant presented an uniform cell morphology without the presence of large Titan-like cells (Figure 5C). These results indicate that Aqp1 contributes to cell body expansion in response to low-inoculum growth in capsule-inducing conditions, with its absence limiting size and its overexpression enhancing cellular and capsular size.

**Figure 5.**
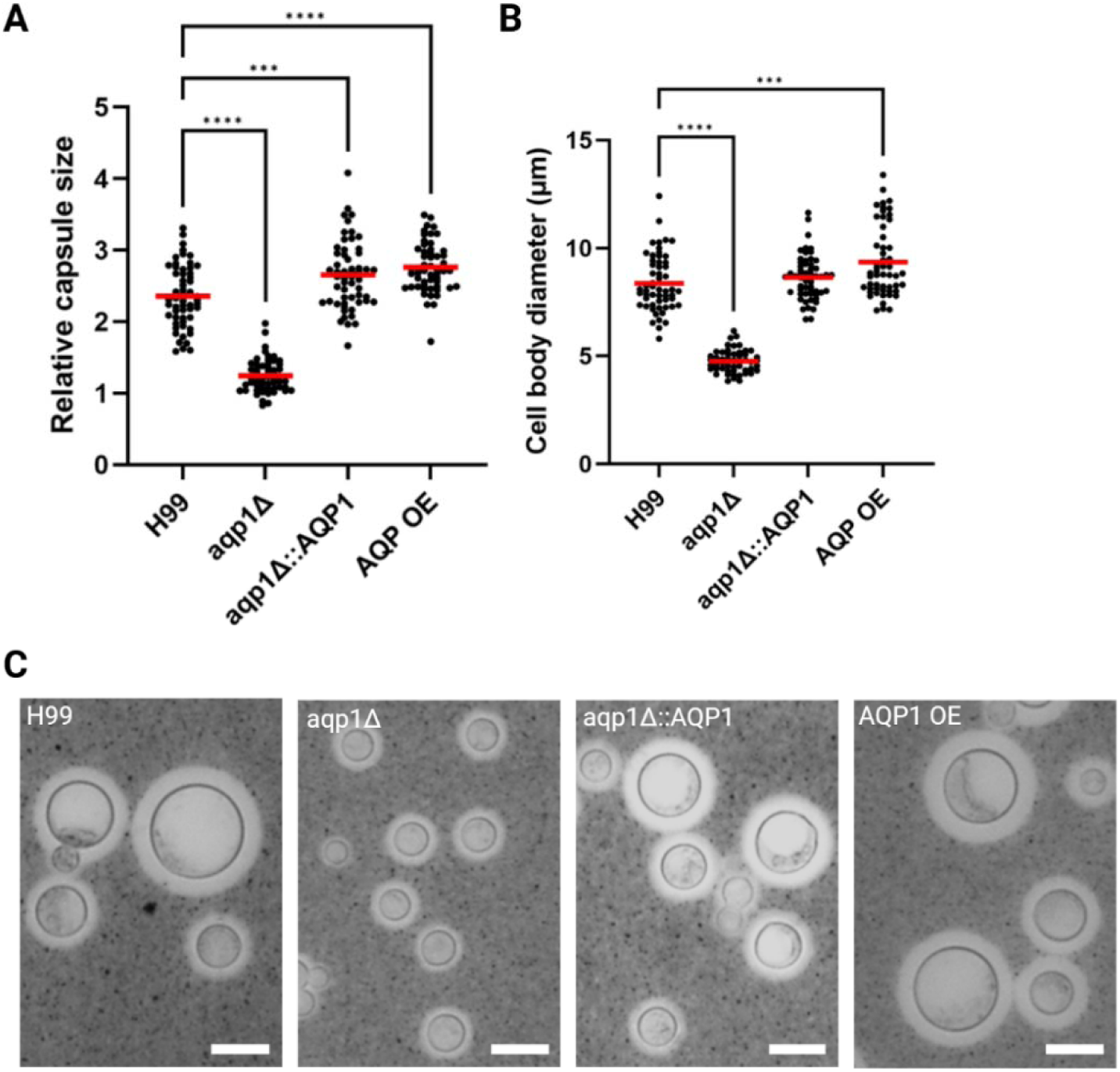
Analysis of cellular morphology in wild-type, AQP1 deletion, complemented, and overexpression strains grown from low inoculum in minimal medium. (A) Scatter plot showing relative capsule size across the four strains. (B) Scatter plot showing cell body diameter measurements for the same strains. Each point represents an individual cell; bars indicate the mean. (C) Representative India ink-stained images of each strain captured under a 40× objective, illustrating differences in capsule and cell body size. Scale bar represents 10lllμm. Data represents the mean from three independent biological replicates. Statistical analysis was performed using 2way ANOVA test.

### Aqp1-Mediated Cell Enlargement Is pH-Dependent but Uncoupled from Growth Rate Changes in *C. neoformans*

A standard Minimal media used to induce the production of cryptococcal polysaccharide capsule was adjusted to acidic pH 5.5. To evaluate if Aqp1-associated cell enlargement could be influenced by extracellular pH, we cultured the wild-type and AQP1-overexpressing strains in MM adjusted to pH 4.5, 5.5, 6.5, or neutral pH 7. After 3 days of incubation at 30 °C, cells were suspended in India ink and analyzed by light microscopy to quantify cell size. At all tested pH levels, the AQP1-overexpressing mutant produced significantly larger cells than the wild-type strain (Figure 6A and 6B).

**Figure 6.**
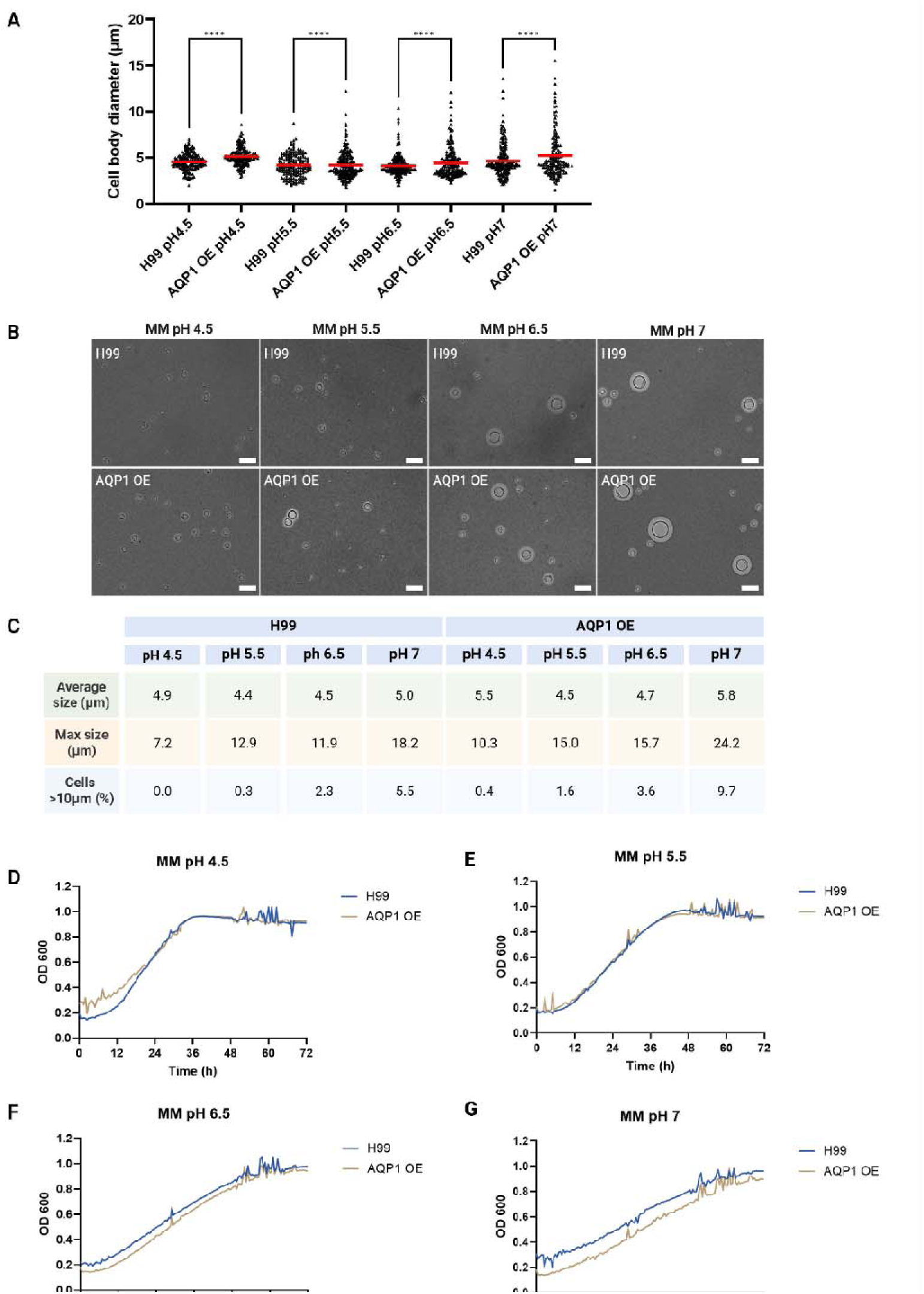
Effect of pH on cell morphology and growth dynamics in wild-type and AQP1-overexpressing (AQP1 OE) strains. (A) Scatter plot showing cell body diameter of wild-type (H99) and AQP1 OE strains grown in minimal medium at pH 4.5, 5.5, 6.5, and 7.0. Each point represents an individual cell; red bars indicate the mean. (B) Representative India ink–stained images captured under a 40× objective for each strain and pH condition, illustrating morphological differences. (C) Summary table presenting the average and maximal cell body diameter for both strains at each pH, along with the percentage of the cell population exceeding 10 µm in diameter. Growth curves of wild-type and AQP1 OE strains in minimal medium adjusted to pH 4.5 (D), 5.5 (E), 6.5 (F), and 7.0 (G). Optical density at 600 nm was measured over time to assess growth dynamics. Data in panel A, D, E, F and G represents the mean from three independent biological replicates. Scale bar = 10 µm. Statistical analysis was performed using 2way ANOVA test.

However, maximal cell sizes in both strains decreased as pH decreased. Specifically, in media at pH 4.5, the largest observed cell diameters were 7.2 µm for wild-type and 10.3 µm for AQP1 OE, whereas at pH 7.0 they reached 18.2 µm and 24.2 µm for wild-type and aquaporin overexpressing mutant strains, respectively. Additionally, the proportion of cells exceeding 10 µm in diameter increased with pH in both strains, ranging from 0% at pH 4.5 to 5.5% (WT) and 9.7% (AQP1 OE) at pH 7 (Figure 6C). These findings suggest that Aqp1-mediated cell enlargement is pH-sensitive, with more pronounced effects at neutral pH.

After observing that cellular size was pH-dependent in an aquaporin-dependent manner, we analyzed whether AQP1 could influence cellular growth rate under varying pH conditions. In this experiment, we compared the growth curves of wild-type and AQP1-overexpressing strains in minimal media adjusted to pH 4.5, 5.5, 6.5, and 7.0. No measurable differences in growth rates were observed between the two strains at pH 4.5, 5.5, or 6.5 (Figure 6D, E, and F). At pH 7.0, the wild-type strain proliferation rate was slightly faster than in the case of the overexpressing mutant strain; however, the difference was minimal and unlikely to be biologically significant (Figure 6G). These results indicate that AQP1 overexpression did not substantially impact growth under standard minimal media conditions across a range of different pH values. However, increasing the pH of the MM had a pronounced adverse effect on overall growth, greatly extending the time required for cultures to reach the plateau phase, suggesting that growth was slowed under less acidic conditions.

### Aquaporin modulated Ion content in *C. neoformans*

To explore the involvement of *AQP1* in maintaining ion balance in *C. neoformans* cells, we utilized Inductively Coupled Plasma Mass Spectrometry (ICP-MS) on wild-type, *aqp1*Δ, complemented, and overexpression strains grown in MM. (Figure 7). The aquaporin deletion mutant presented a significantly lower intracellular potassium (K) concentration than the wild type (Figure 7A). In contrast, both the complemented and overexpression strains showed slightly elevated K levels compared with the wild-type strain. All three mutant strains displayed slightly higher intracellular phosphorus (P) concentrations than the WT (Figure 8B). Notably, the deletion mutant presented substantially elevated levels of copper (Cu) and zinc (Zn) compared to wild type, suggesting a potential role for AQP1 in maintaining metal ion balance (Figure 7F and 7G). These data indicate a link between AQP1 expression and the regulation of specific intracellular element concentrations. Given that the *AQP1* deletion mutant showed significantly elevated intracellular levels of copper (Cu) and zinc (Zn) compared to the wild-type strain, we hypothesized that *AQP1* may influence the function of copper-and zinc-dependent enzymes such as superoxide dismutase 1 (SOD1). We thus examined Cu and Zn tolerance in H99 and aquaporin mutant strains. The AQP1 OE strain grew slightly better at 0.312 mM CuSO at 24 h, but by 48–72 h, all strains showed similar growth at 0.625 mM CuSO (Supplementary figure 3 A–C). Zinc tolerance was comparable across strains, with growth at 1.25 mM ZnSO after 48–72 h (Supplementary figure 3 D–F). These results indicate that aquaporin expression has little effect on metal resistance in *C. neoformans*.

**Figure 7.**
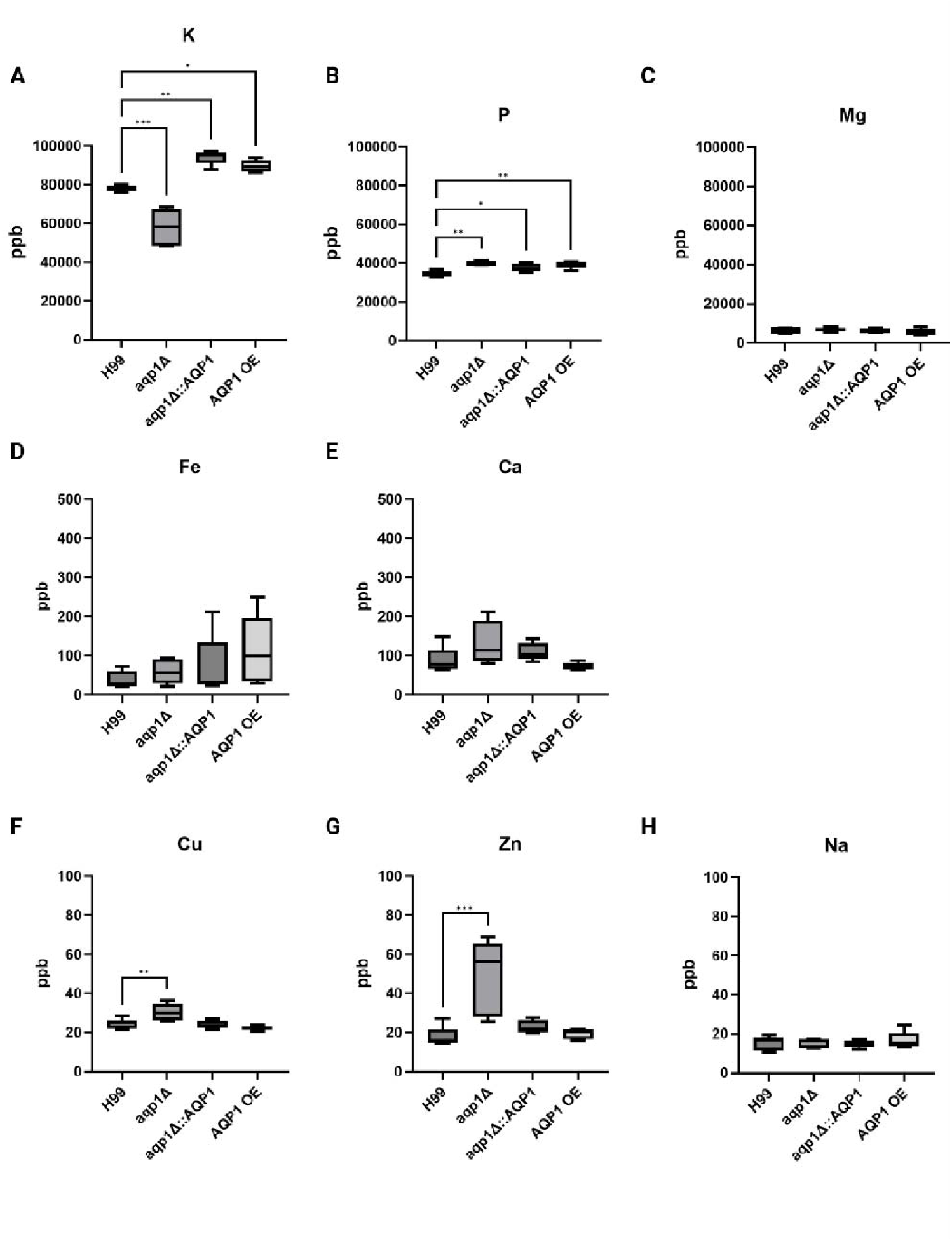
Inductively Coupled Plasma Mass Spectrometry (ICP-MS) analysis of selected elemental concentrations in wild-type, AQP1 deletion, complemented, and overexpression strains grown in standard minimal media. Box plots represent the mean from 5 biological replicates. Elements analyzed include (A) potassium (K), (B) phosphorus (P), (C) magnesium (Mg), (D) iron (Fe), (E) calcium (Ca), (F) copper (Cu), (G) zinc (Zn), and (H) sodium (Na). Elemental concentrations were assessed to evaluate potential roles of AQP1 in ion homeostasis and transport. Data represents the mean of five independent biological replicates. Statistical analysis was performed using 2way ANOVA test.

**Figure 8.**
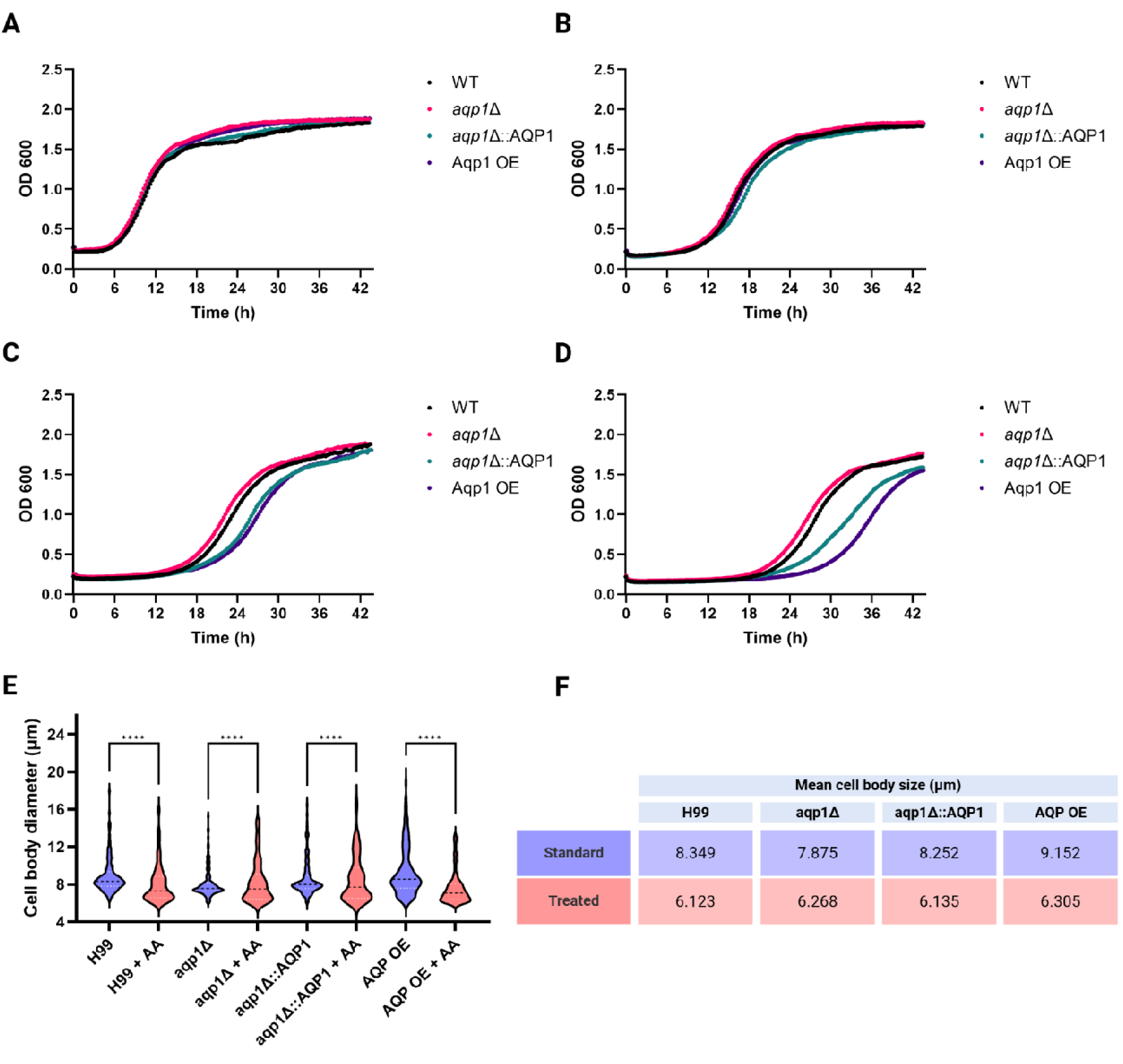
Impact of ROS and antioxidant treatment on growth and cell morphology in Cryptococcus neoformans. Growth dynamics and cell size distribution were analyzed in four C. neoformans strains: wild-type H99, aqp1Δ, aqp1Δ::AQP1, and AQP1 OE. Panels A–D show growth curves in YPD medium under increasing oxidative stress induced by menadione: (A) control (YPD only); (B) YPD + 25 µM menadione (C) YPD + 50 µM menadione; (D) YPD + 75 µM menadione. (E) violin plots of cell body sizes (>6 µm diameter) under titan cell-inducing conditions, comparing untreated and 3 mM ascorbic acid (AA)-treated cultures. (F) table summarizing the data from panel E, showing average cell sizes for each strain with and without AA treatment. Data represents the mean from three independent biological replicates. Statistical analysis was performed using 2way ANOVA test.

### Cryptococcal aquaporin modulates growth and intracellular ROS homeostasis

To examine if AQP1 plays a role in the oxidative stress response, we examined growth dynamics and cell morphology under conditions of reactive oxygen species (ROS) stress and antioxidant treatment (Figure 8). Growth curves were generated for WT, *and a set* of aquaporin mutant strains cultured in YPD medium with increasing concentrations (0, 25, 50, and 75 µM) of menadione (vitamin K3), which promotes oxidative damage. While all strains grew similarly under control conditions (0 µM), the *AQP1*-overexpressing strain exhibited a reduced growth rate relative to the wild-type and deletion strains when exposed to 50 and 75 µM menadione (Figure 8 A–D), suggesting that *AQP1* overexpression increased sensitivity to ROS stress. Visualization of intracellular ROS levels revealed subtle differences in fluorescent signal intensity among the tested strains (Supplementary figure 4). The wild type, *aqp1*Δ mutant, and complemented strains showed comparable levels of intracellular staining, whereas the AQP1 overexpression strain exhibited a consistently brighter fluorescence profile, indicating greater ROS accumulation within the cells. These differences further support the idea that AQP1 overexpression leads to accumulation of intracellular ROS levels in *C. neoformans* cells.

### Aqp1 localizes to intracellular membranes and modulates titan cell formation and vacuole morphology during *In vitro* induction

As reported in previous research, aquaporin 1 localizes to the internal membrane in yeast cells of *C. neoformans* (Figure 9A)(12). Visualization of aquaporin localization in the titan cells revealed its presence on the intracellular membrane at the periphery of the titan cells, in the cytoplasmic compartment of cells surrounding the central large vacuole (Figure 9B).

**Figure 9.**
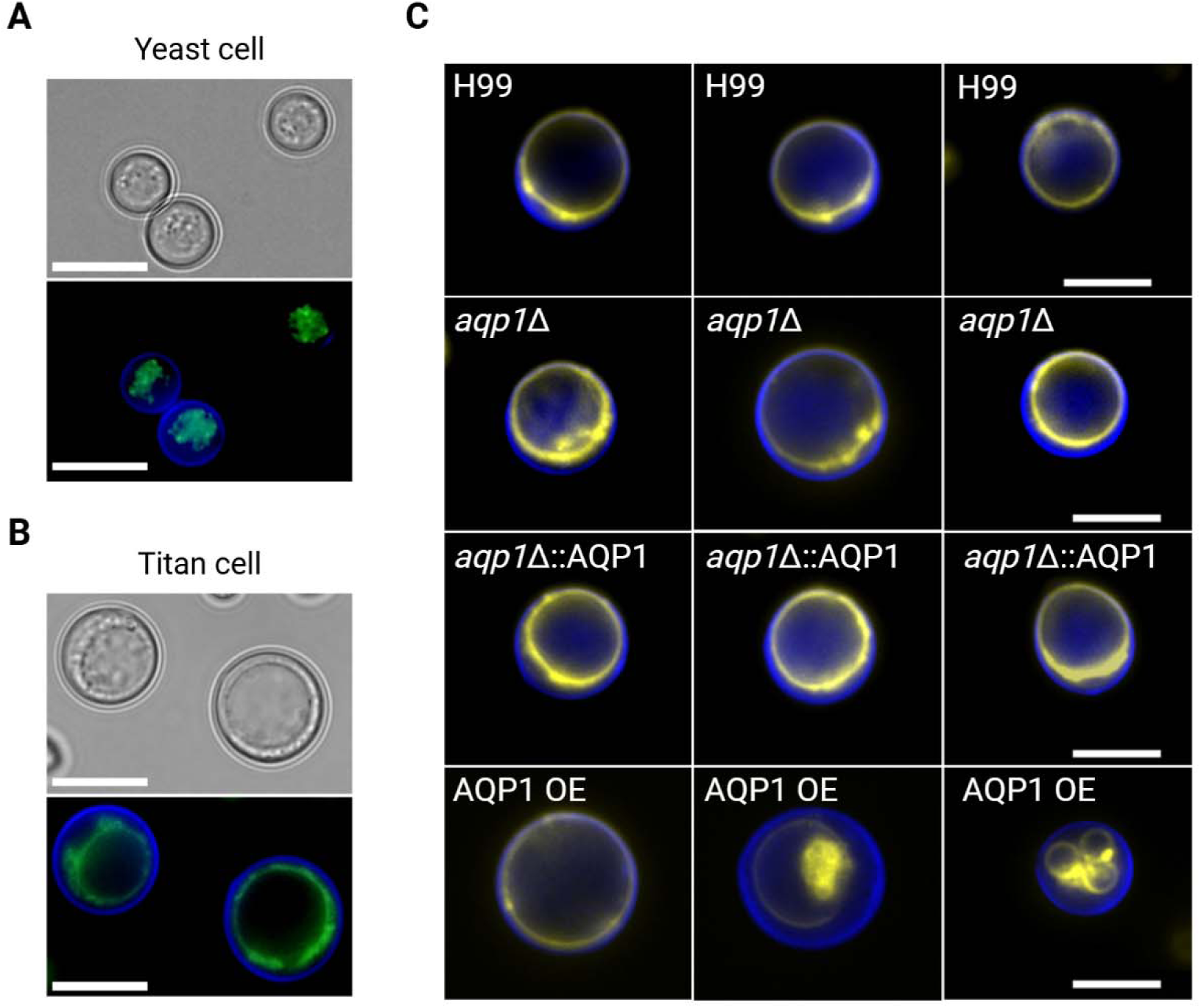
Localization of Aqp1 and analysis of vacuole morphology in yeast and titan-like cells. (A) Differential interference contrast (DIC) and fluorescence images of yeast-form C. neoformans cells expressing Aqp1-GFP. Green fluorescence indicates Aqp1-GFP localization, and the cell wall was stained with Uvitex 2B (blue). (B) DIC and fluorescence images of titan-like cells expressing Aqp1-GFP under titan cell–inducing conditions. Green fluorescence indicates the localization of GFP-tagged aquaporin in titan-like cells, and the cell wall was stained with Uvitex 2B (blue). (C) Analysis of vacuole morphology in titan-like cells of WT, aqp1Δ, aqp1Δ::AQP1, and AQP1 OE strains. Vacuoles were stained with FM4-64 (yellow), and cell walls were stained with Uvitex 2B (blue). Scale bar = 10 µm.

Given our earlier observations of frequent cell enlargement in the AQP1-overexpressing strain, we investigated whether aquaporin 1 influenced the intracellular morphology of titan cells in *C. neoformans*. One of the significant characteristics of cryptococcal titan cells is their large central vacuole (31, 33). Vacuolar FM4-64 staining revealed striking vacuolar abnormalities in vacuolar morphology in the AQP1-overexpressing strain affecting approximately 24% of titan-like enlarged cells (Figure 9C). In contrast, the frequency of vacuolar defects in titan-like cells of the wild-type, *aqp1*Δ, and complemented strains remained close to 0%. Together, these findings suggest that Aqp1 facilitates titan cell enlargement and that its overexpression may disrupt vacuole homeostasis in titan-like cells.

## Discussion

Aquaporins are integral membrane proteins known for functioning to facilitate water transport across biological membranes, yet their roles in fungal pathogens, particularly *Cryptococcus neoformans*, remain incompletely understood. Prior to this study, Aqp1 in *C. neoformans* was known to be transcriptionally induced by osmotic and oxidative stress via the HOG pathway yet was largely dispensable for adaptation to environmental stressors and virulence (12). Aqp1 was shown to play a minor role in regulating cell-surface hydrophobicity and to be involved in maintaining metabolic homeostasis (12). Deletion of AQP1 led to the accumulation of various metabolites and enhanced competitive survival, while overexpression of AQP1 reduced metabolite levels (12). Here, we initially sought to evaluate potential contribution of aquaporin in the process of thermal stress response and to test if aquaporin channel plays an important role in regulation of cellular buoyancy. In this study, we investigated the contribution of the *C. neoformans* aquaporin Aqp1 to growth, morphogenesis, and titan cell formation under various environmental conditions. Our results reveal that Aqp1 does not influence cryptococcal growth under standard, nutrient-rich laboratory culture conditions, but significantly impacts cell enlargement and morphological adaptation in host-like environments that confer cellular stress and may additionally have a role in protection against heavy metals.

The growth analyses presented here support a role for Aqp1 in glycerol utilization by *C. neoformans*. Strains expressing Aqp1 grew normally in media containing glucose, indicating that this protein was not required for metabolism of the preferred carbon source. However, clear differences emerged when glycerol was provided as the main carbon source. Under those conditions, cells lacking AQP1 showed reduced proliferation, while cells overexpressing the gene showed increased growth. These findings support previous predictions that Aqp1 acts as an aquaglyceroporin, facilitating the movement of glycerol across the cell membrane. Initial phenotypic analyses using spot assays demonstrated that neither deletion nor overexpression of *AQP1* altered growth under thermal stress (30°C, 37°C, or 39°C) or during prolonged incubation at low temperatures (4°C). Elimination or overexpression of aquaporin in C*. neoformans* had no visible effect on its susceptibility to fluconazole treatment. Similarly, Aqp1 did not affect cell size, capsule thickness, or buoyancy in nutrient-rich medium, suggesting that under non-capsule-inducing conditions, Aqp1 was dispensable for basic cellular physiology. In contrast, under capsule-inducing conditions, particularly in minimal medium, Aqp1 overexpression resulted in significantly increased capsule size and cell body diameter, with a notable subpopulation of cells exceeding 10 µm in diameter—reminiscent of titan cells (32–34, 50–52). This phenotype was further accentuated when cells were cultured using in low density conditions, which are known to favor morphological diversification. The *aqp1* deletion mutant exhibited significantly reduced cell size, whereas overexpression of Aqp1 maintained an enlarged phenotype, supporting a role for Aqp1 in promoting cellular expansion. Since pH can modulate fungal physiology, including capsule formation and cell size, we investigated whether Aqp1-mediated cell enlargement is correlated with a different spectrum of pH (53–55). Our findings show that less acidic pH conditions promoted the emergence of enlarged and titan-like cells in both wild-type and AQP1-overexpressing strains, with a pH-dependent increase in the proportion of cells >10 µm, which is consistent with previous observations suggesting the importance of neutral pH on titan cells formation (31).

Interestingly, the increase in pH not only favored the emergence of enlarged and titan-like cells but also correlated with a slower overall growth rate, independently of AQP1 expression. Cultures grown at higher, less acidic pH levels exhibited delayed progression to the plateau phase, suggesting that more neutral conditions may impose physiological constraints that decelerate proliferation while promoting morphological remodeling.

Using established *in vitro* titan cell induction media, we tested the role of Aqp1 in titan cell formation. Titan cells are critical for the virulence of *C. neoformans* because their enlarged size makes them resistant to phagocytosis by host immune cells, enhancing survival within the host (33, 56). Formation of the titan cell during the infection may contribute to immune modulation and promote dissemination by generating smaller, progeny cells (27, 33, 35). The overexpression of *AQP1* led to a significant increase in average cell size and frequency of titan-like cells, whereas the *aqp1*Δ mutant failed to undergo typical size enlargement.

Here, we also show that Aqp1 is involved in intracellular ionic, oxidative, and morphological homeostasis in *C. neoformans*. The deletion mutant showed significantly reduced intracellular levels of potassium, copper, and zinc, indicating that Aqp1 can influence ion balance. These findings suggest that cryptococcal aquaporin may play a key role in metal distribution and metabolism, comparable to the role of mammalian Aqp4 in the brain (57). The elevated Cu and Zn levels are particularly notable given their essential role as cofactors for superoxide dismutase 1 (Sod1), a major antioxidant enzyme that protects against various forms of reactive oxygen species. Both metals are also toxic to fungi at higher concentrations (58, 59) (60). To explore this connection, we analyzed the response of AQP1-altered strains to oxidative stress and found that AQP1 overexpression increased sensitivity to ROS, while its deletion had a modest protective effect. Additionally, titan cell formation, which is linked to virulence and immune evasion, was modulated not only by ROS levels but also by AQP1 expression (61). The AQP1-overexpressing strain exhibited the largest titan-like cells, while the *aqp1*Δ strain formed a population of significantly smaller cells. In addition, antioxidant supplementation suppressed cell enlargement, confirming previously observed roles for oxidative signaling in titan morphogenesis (61). These data support a model in which Aqp1 influences not only water and solute flux but also intracellular redox balance, potentially through its effects on metal ion availability and Sod1 activity, thereby shaping the pathogen’s stress response and morphological adaptation. One defining feature of cryptococcal titan cells is the presence of a large vacuole, which is involved in maintaining intracellular organization and turgor pressure (31, 33, 62, 63). In our study, *AQP1* overexpression led to frequent vacuolar abnormalities in titan-like cells, a phenotype rarely seen in wild-type or control strains. This suggests that while proper activity of Aqp1 promotes cell enlargement, its overexpression may disrupt vacuole integrity, possibly because excessive water influx overwhelms the cell’s ability to maintain appropriate vacuolar morphology.

Taken together, our findings establish a novel role for the aquaporin in regulating cell enlargement, stress adaptation, and virulence factor expression in *C. neoformans*.

While it is not essential for growth under optimal conditions, Aqp1 is a key factor for navigating morphogenetic responses to host-like stimuli such as low nutrient availability, pH variation, or oxidative stress. In this study, we demonstrate that cryptococcal Aqp1 influences intracellular ion composition, including levels of elements essential for antioxidant defense. Its presence can modulate the intracellular response to ROS, which in turn can affect the formation of titan cells. Finally, AQP1 overexpression leads to vacuolar abnormalities in titan-like cells, suggesting that unbalanced water influx may compromise intracellular organization during enhanced cell expansion. Overall, our findings provide new insights into molecular function of a fungal aquaporin with specific focus on ROS-dependent morphogenesis of cryptococcal titan cells..

## Experimental procedures

### Strains and media

Wild-type *C. neoformans* strain H99 and all aquaporin deletion, complemented strain, and overexpression mutants were obtained from the Yun Sun Bahn Laboratory (Supplementary Table 1) at Yonsei University (South Korea) (12). All strains were routinely cultured in YPD liquid or solid medium (BD Difco, Sparks, USA) at 30 °C.

### Structural modeling and functional prediction

The similarity of the amino acid primary sequence of cryptococcal Aqp1 (CNAG_01742) was compared to human aquaporins using BLASTp. To gain insight into the structural characteristics of the cryptococcal aquaporin Aqp1, we performed 3D protein structure prediction using AlphaFold3 (64) Similar prediction structural analysis was made for Aqp1 homologues present in other *Cryptococcus gattii*, *Malassezia furfur*, *Candidozyma auris*, *Candida parapsilosis* and *Mucor circinelloides*. AlphaFold structures of cryptococcal Aqp1 and human AQP1 and AQP8 were structurally aligned with a Smith-Waterman algorithm using UCSF ChimeraX version 1.9 (65). The DeepTMHMM topology prediction model and SignalP 6.0 signal peptide prediction algorithm were used to evaluate the additional C-terminal domain of the cryptococcal Aqp1 (65, 66). The search for aquaporin domain (PF00230) containing proteins in other clinically relevant fungal species was performed via OrthoMCL DB (67).

### Serial spot dilution assay for thermal and fluconazole sensitivity

Overnight YPD cultures were washed with DPBS, adjusted to a final concentration of 10^4^ cells in 3 µL, and used for a 10-fold serial dilution. Cells were spotted onto solid media in 3 μL spots, and the plates were incubated at the respective temperature and photographed. To assess the melanization of cryptococcal cells, overnight cultures were washed with DPBS, adjusted to a final concentration of 10^4^. A total of 3 μL of each culture were spotted onto MM agar media supplemented with L-DOPA and incubated for 3 d at 30°C.

### Glycerol-based growth analysis

Yeast Nitrogen Base w/o amino acids and ammonium sulfate (YNB) (Difco) medium was prepared by dissolving 1.7 g of YNB base and 40 mM (NH) SO (Sigma) in 1 L of water and supplementing with 111 mM of the indicated carbon source (glucose or glycerol). The pH was adjusted to 5.5 before sterilization with a vacuum filtration system Stericup® (MilliporeSigma). Minimal media (MM) was prepared with 111 mM of carbon source (glucose or glycerol), 13 mM glycine (Sigma), 10 mM MgSO (Sigma), 29.4 mM KH PO (Sigma), 3 µM thiamine-HCl, and adjusted to pH 5.5. A single colony of each strain was used to inoculate 4 mL of YPD media in 14 mL bottom-round tubes (Falcon), for 2d with agitation at 30 °C. Yeast cells were harvested, washed 2 times with DPBS, and counted with a hemocytometer. Subsequently, 1×10^6^ cells/mL were inoculated into 14 mL bottom-round tubes containing 4 mL of YNB or MM media with glycerol or glucose as the sole carbon source and incubated with agitation at 30 °C. Glucose media were used as a control. The cells were counted daily throughout the experiment.

Results represent the cell density in the entire sample. Growth data were visualized and analyzed using GraphPad Prism 10.1.1.

### Cell morphology, buoyancy, and hydrophobicity assay

Cell cultures were incubated in YPD liquid media for 2 days at 30 °C and washed with DPBS. To evaluate cell diameter and capsule size, cells were stained with India ink and photographed on an Olympus AX70 microscope (Olympus America, USA). Cell size was established based on measurement of cell diameter, and relative capsule size was established by measuring the capsule diameter to cell diameter ratio. To assess buoyancy, 3 mL of stationary-phase YPD liquid cultures for each strain were pipetted into cuvettes and allowed to settle passively for 6 hours. Photographs were taken hourly for each cuvette, and the displacement from the top of the cuvette was measured to generate a graph of the displacement rate over time, as previously described (68). To measure cell surface hydrophobicity, wild-type and aquaporin mutant cultures were analyzed using the microbial adhesion to hydrocarbons (MATH) assay as previously described (69). In short, cells were grown overnight in YPD liquid media for 2 days at 30 °C and washed with DPBS. Cell cultures were suspended in 3ml of DPBS with an OD600 ∼0.3, mixed with 400 µL hexadecane (Sigma-Aldrich), vortexed for 1 min, and rested for 2 min. After the settlement period, the hydrophobic layer and the bottom layer (aqueous layer) were transferred to a 96-well plate, and the OD600 was measured in the SpectraMAX 340 Tunable Microplate Reader (Molecular Devices Ltd).

### Analysis of cell body and capsule size in capsule inducing minimal media

To induce polysaccharide capsule growth, a 2 d YPD culture was washed with DPBS, and 10 µL of culture was transferred into 5 mL of minimal media (7.5 mM glucose, 10 mM MgSO4, 29.4 mM KH2PO4, 6.5 mM glycine, and 3 µM thiamine-HCl, pH 5.5) and incubated for 3 d at 30°C, on the Cel gyro Rotator (Thermo Scientific). For long term analysis cells were incubated in 5 mL of minimal media and 100 µL of tested cultures were taken every two days to visualize cell and capsule size.

Alternative versions of minimal media were prepared at pH 4.5, 5.5, 6.5, and 7. To evaluate cell diameter and capsule size, cells were stained with India ink and photographed on an Olympus AX70 microscope. Cell and capsule size were measured using ImageJ and plotted using GraphPad Prism 10.1.1.

### Growth curve analysis in minimal media at varying pH

To assess the effect of AQP1 overexpression on pH-dependent growth, H99 and AQP1-overexpressing *C. neoformans* strains were cultured in minimal medium adjusted to different pH levels (pH 4.5, 5.5, 6.5, and 7.0). A 1 volume mL of media was distributed to 24-well plates and inoculated with 5 × 10^5^ cells /m. Cultures were incubated at 25 °C with constant shaking. Optical density at 600 nm (OD) was measured every 30 min over time using a microplate reader (Spectramax M5) to monitor growth kinetics.

Each condition was tested in three independent biological replicates, and results were expressed as the mean OD values over time. Growth data were visualized and analyzed using GraphPad Prism 10.1.1.

### Cellular and Population Level Total Reactive Oxygen Species Analysis

To determine the total ROS between H99, *aqp1*Δ, *aqp1*Δ::AQP1, and AQP1 overexpression strains, overnight YPD cultures were washed twice with DPBS, then cells were treated with 50µM menadione to generate superoxide stress for 3 hours in DPBS in a rotary tube mixer. Cell cultures were washed and stained for 30 min at 37°C with either 10µM dihydroethidium (DHE, ThermoFisher Scientific D23107) for cellular analysis of total ROS or 5uM CellROX (Invitrogen, C10444) for population analysis of total ROS. After two washes, DHE-stained cells were mounted on agarose patch and imaged on Olympus AX70 microscope.

### Titan cells formation and intracellular visualization of Aqp1-GFP and vacuoles

Cells from a 2 d YPD culture were washed with DPBS. The cells were inoculated in titan stimulation medium or on standard MM using a low inoculum. For the first method, 1 x 10^5^ cells/mL were inoculated in a 24-well plate with 1 mL of DPBS with 10% Fetal bovine serum (FBS) and incubated for 72 h at 37°C and 5% CO_2_ (32, 70). For the low-inoculum titan cell induction method, 1×10^3^ cells/mL were inoculated into 5 mL of MM in 14 mL bottom-round tubes for 4 days with agitation. The cells were then recovered, washed with DPBS, and stained with India Ink. To visualize the intracellular localization of cryptococcal aquaporin1, a 2 d YPD culture of the GPF-tagged aquaporin strain was washed with DPBS. Then, 1 x 10^5^ cells/mL were inoculated in a 24-well plate with 10% Fetal bovine serum (FBS) and incubated for 72 h at 37°C and 5% CO_2_ to stimulate titan cell morphology, or cells were incubated for 72 h at 30°C in MM to present standard size yeast cells. After incubation, the cells were washed twice with DPBS and observed using an Olympus AX70 microscope. Cell size for titan-like cells was measured using ImageJ and plotted using GraphPad Prism 10.1.1.

For vacuolar and cell wall staining, cell cultures were labeled with FM4-64 (1 μg/mL) and UVitex 2B (5 μg/mL) for 30 min at 30 °C. After incubation, the cells were washed twice with DPBS and observed using an Olympus AX70 microscope. Cell size for titan-like cells was measured using ImageJ and plotted using GraphPad Prism 10.1.1.

### Inductively Coupled Plasma Mass Spectrometry

Selected cryptococcal cultures were incubated in 5 mL of minimal media in the rotating wheel for 3 days. Cell cultures were pelleted and washed twice with TE buffer (10 mM Tris, 1 mM EDTA, pH 8.0), then twice with MiliQ H_2_O. Washed cell pellets were lyophilized to remove excess liquid and digested in 20% HNO_3_ overnight at 85°C in metal-free 15 mL conical tubes. Digest was diluted to 5% HNO3 and analyzed at the Molecular Characterization and Analysis Complex at the University of Baltimore County for Fe, Mn, Zn, Cu, Ca, K, Na, P, and Mg analysis via PerkinElmer NexION 300D with ICP.

### Growth analysis in the presence of copper and zinc

To evaluate the impact of Cu or Zn on growth, *C. neoformans* strains (H99, *aqp1*Δ, *aqp1*Δ::*AQP1*, and *AQP1* overexpression) were cultured in minimal medium with varying concentration of CuSO_4_ (5 to 0.156 mM) (Sigma) or ZnSO_4_ (5 to 0.156 mM) (Sigma). Growth assays were performed in a 96 wells microtiter plate with a biological triplicate, with continuous agitation at 30 °C, and optical density at 600 nm (OD) was measured using a microplate reader (Spectramax M5). Growth curves plots were visualized using GraphPad Prism 10.1.1.

### Assessment of growth under oxidative stress and antioxidant treatment

To analyze if oxidative stress may have a stimulative impact on growth rate, *C. neoformans* strains (H99, *aqp1*Δ, *aqp1*Δ*::AQP1*, and *AQP1* overexpression) were cultured in YPD liquid medium supplemented with different concentrations of menadione (0, 25, 50, and 75 µM). Growth assays were performed in microtiter plates with continuous shaking at 30 °C, and optical density at 600 nm (OD) was measured every 30 min using a microplate reader (Spectramax M5) to generate growth curves.

Growth curves plots were visualized using GraphPad Prism 10.1.1.

Evaluation of antioxidant supplementation on the formation of titan cells was performed in titan cell-inducing conditions. In short, cryptococcal cells were incubated for 72 h in titan-inducing conditions (PBS with 10% heat-inactivated FBS, 37°C, 5% CO_2_), with or without 3 mM ascorbic acid (AA), to assess if antioxidants can limit aquaporin-associated increases in cell size. After incubation, cell body diameters were measured under a microscope (Olympus AX70) using ImageJ software. Only cells exceeding 6 µm in diameter were included in the analysis. Cell size distributions were visualized using GraphPad Prism 10.1.1.

### Cellular and Population Level Total Reactive Oxygen Species Analysis

To determine the total ROS between H99, *aqp1*Δ, *aqp1*Δ*::AQP1*, and *AQP1* overexpression strains, overnight YPD cultures were washed 2x with PBS, and cells were treated with 50 uM menadione to generate superoxide stress for 3 h in PBS in a rotary tube mixer. Cells were washed and stained for 30 min at 37 °C with either 10 μM dihydroethidium (DHE, ThermoFIsher Scientific D23107) for cellular analysis of total ROS or 5 μM CellROX (Invitrogen, C10444) for population analysis of total ROS. After 2x washes in PBS, DHE-stained cells were mounted on an agarose patch and imaged on a Leica THUNDER widefield microscope.

## Acknowledgement

We thank Professor Yun Sun Bahn and his laboratory (Yonsei University, South Korea) for providing the *Cryptococcus neoformans* strain H99 aquaporin **aqp1** deletion mutant, complemented mutant, and overexpression mutant. A.C. was supported in part by the National Institutes of Health (NIH) grants AI052733, AI15207, AI171093-01, and HL059842

