## Supplementary figures and table for "A role for aquaporin (Aqp1) in the control of *Cryptococcus neoformans* cell morphology"

**Supplementary Table 1. List of strains used in this study.**

| **Strain** | **Original name** | **Genotype** | **Parent** |
| --- | --- | --- | --- |
| H99 | H99 | MATα |  |
| *aqp1*Δ | YSB395 | MATα aqp1::NATSTM#191 | H99 |
| *aqp1*Δ::AQP1 | YSB2419 | MATα cnag_01742Δ(APQ1)::NAT+pJAF12_AQP1::NEO | YSB395 |
| AQP1-GFP | YSB3710 | MATα AQP1:GFP::NEO (CNAG_01742) | H99 |
| AQP1 OE | YSB2350 | MATα PH3:AQP1‐NAT (AQP1 overexpression strain) | H99 |

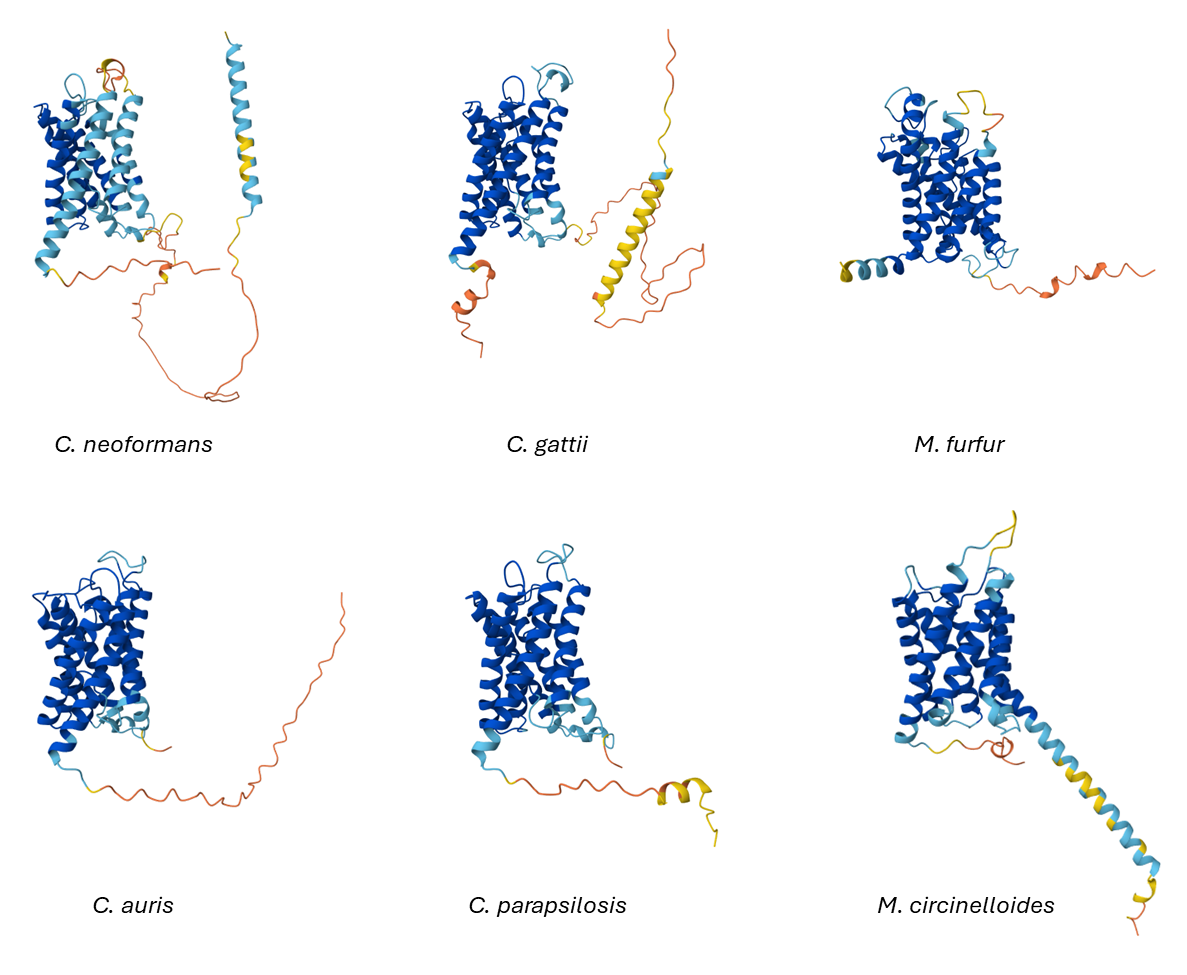

**Supplementary figure 1. Predicted three-dimensional structures of aquaporin homologs from six fungal species generated using AlphaFold3.** Structural models are shown for *Cryptococcus neoformans*, *Cryptococcus gattii*, *Malassezia furfur*, *Candidozyma auris*, *Candida parapsilosis*, and *Mucor circinelloides*. Each model highlights the conserved transmembrane architecture characteristic of fungal aquaporins, including the canonical pore-forming helices. Models are displayed in the same orientation to facilitate comparison of structural conservation across species.

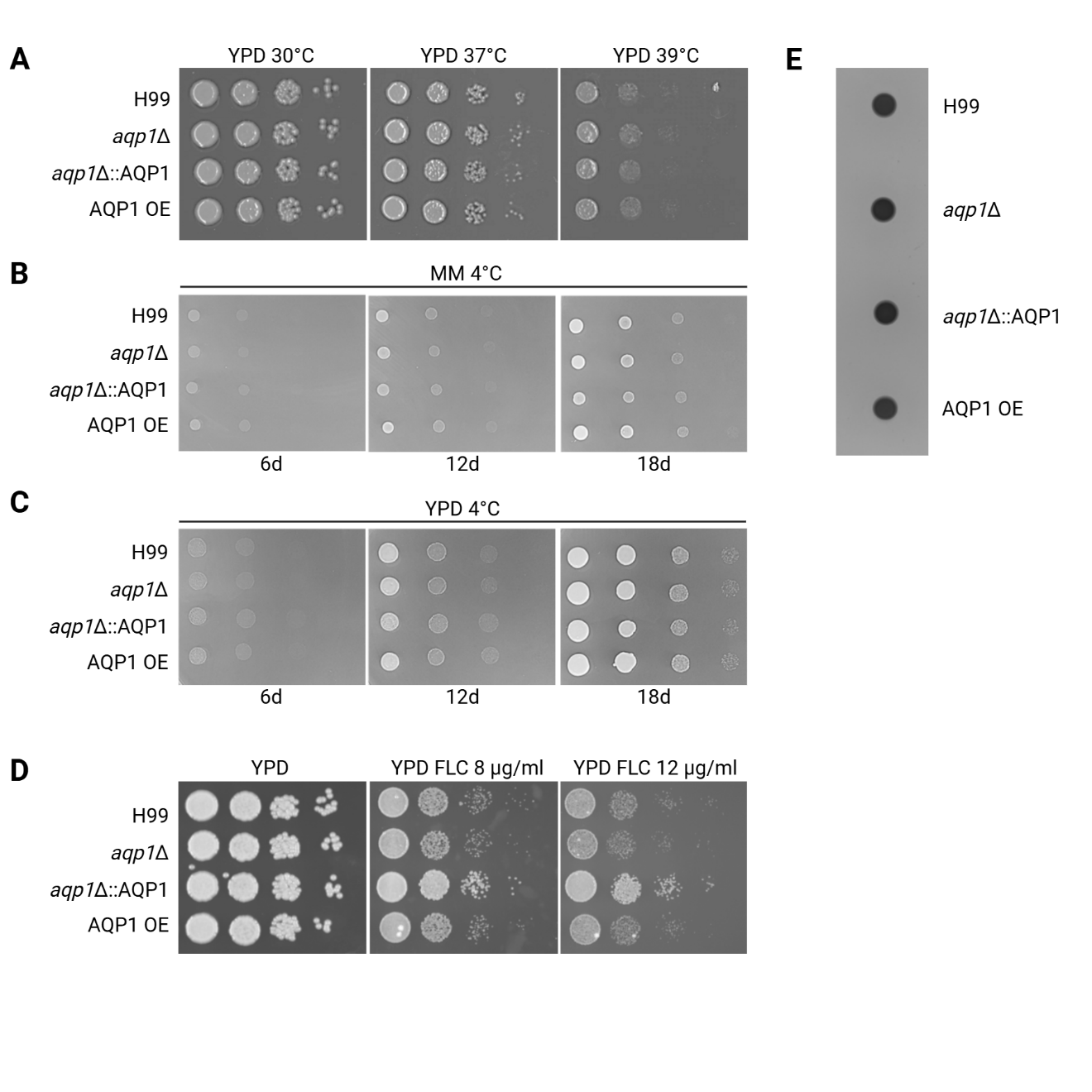

**Supplementary figure 2. Serial dilution assays evaluating the temperature stress response, melanization, and fluconazole sensitivity of wild-type, AQP1 deletion, complemented, and overexpression strains.** (A) Cells were spotted onto YPD plates and incubated at 30 °C, 37 °C, and 39 °C for 2 days to assess growth under heat stress conditions. (B) Cells were spotted onto minimal media plates and incubated at 4 °C to evaluate cold stress tolerance. Colony growth was imaged after 6, 12, and 18 days. (C) Cells were spotted onto YPD plates and incubated at 4 °C to evaluate cold stress tolerance in reach nutrition environment. Colony growth was imaged after 6, 12, and 18 days. (D) Cells were spotted onto YPD media plates with various doses of Fluconazole (FLC) and incubated at 30 °C for 2 days to evaluate the Fluconazole sensitivity of selected strains. (E) Cell cultures of WT and tested mutant strains were spotted onto a solid MM agar plate supplemented with L-DOPA and incubated at 30 °C for 3 days.

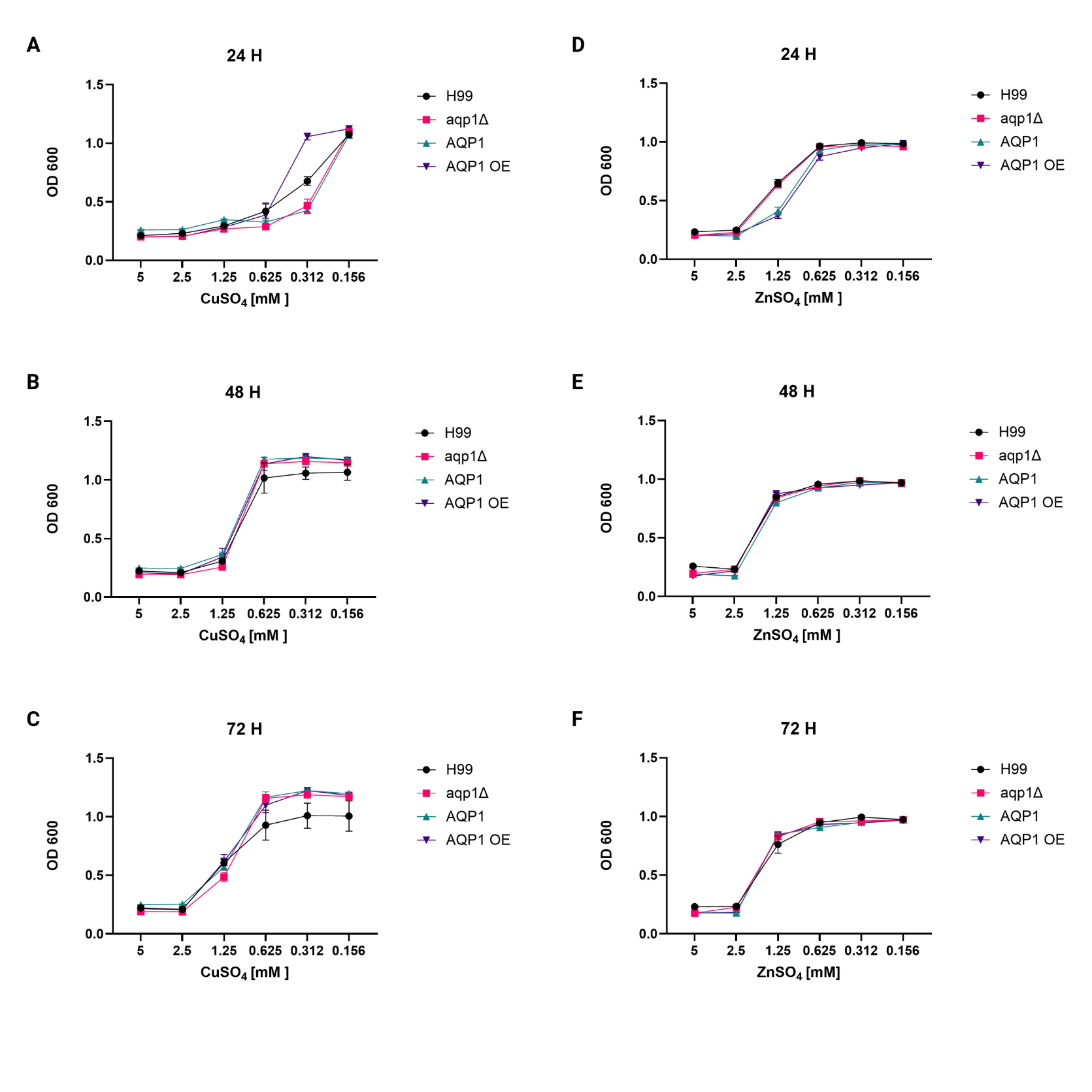

**Supplementary figure 3. The figure presents the growth sensitivity responses of *C. neoformans* strains to Cu and Zn stress.**

Growth dynamic analysis in minimal medium (MM) supplemented with varying concentrations of CuSO_4_ (5 mM to 0.156 mM) was quantified by measuring optical density (OD) at 600 nm. Linear graphs reveal the differential responses of each strain to copper exposure after 24 hours (A), 48 hours (B), and 72 hours (C). Data represents the mean from three independent biological replicates.

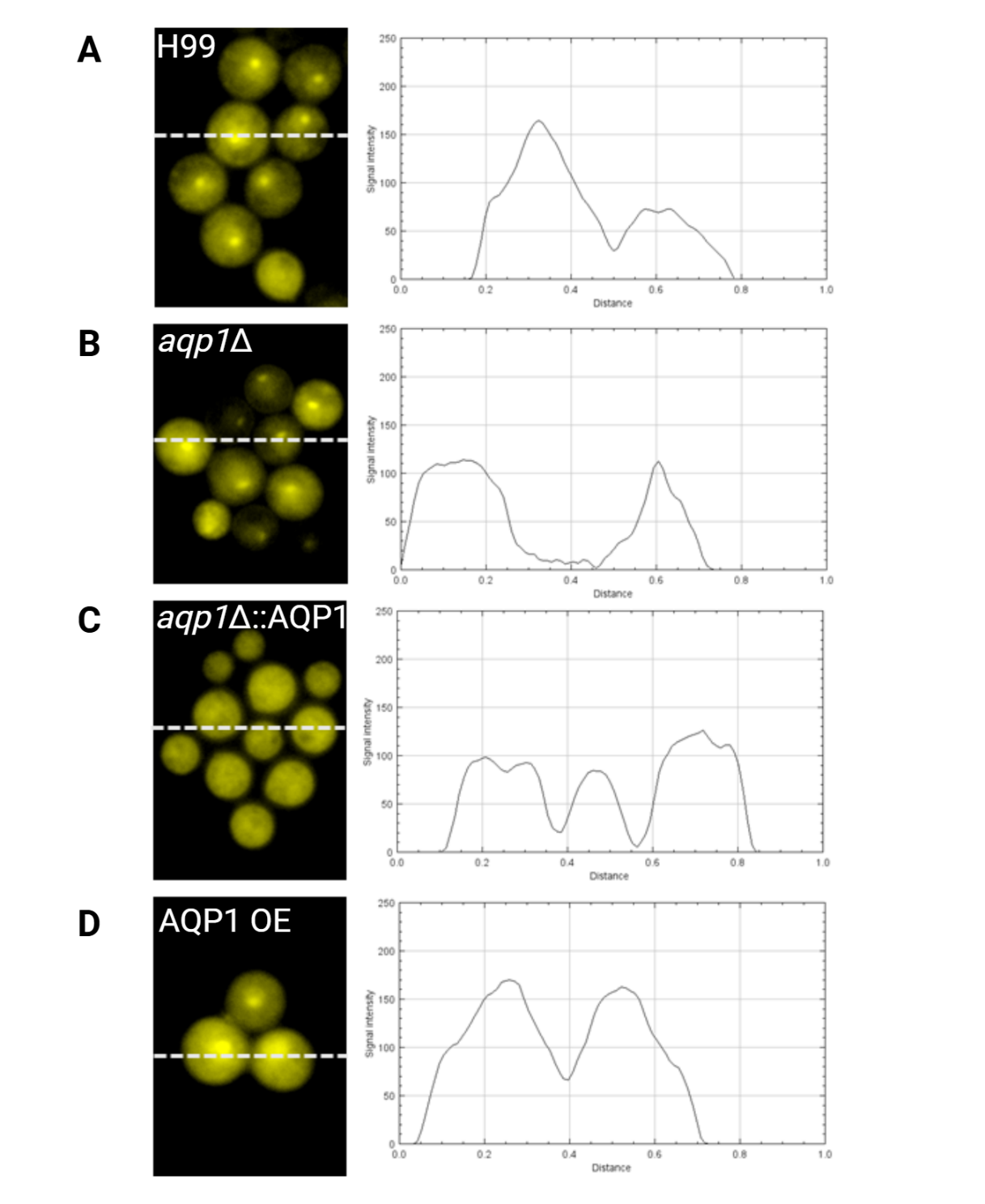

**Supplementary Figure 4. Intracellular reactive oxygen species (ROS) accumulation in *C. neoformans* cells visualized by fluorescence microscopy.** Cells were stained with a ROS-sensitive dye (DHE), and intracellular ROS was visualized as yellow fluorescence. (A) Representative fluorescence image of ROS staining in the wild-type strain H99, accompanied by a corresponding fluorescence-intensity profile. (B) ROS staining and intensity profile for the *aqp1*Δ deletion mutant. (C) ROS staining and intensity profile for the *aqp1*Δ::AQP1 complemented strain. (D) ROS staining and intensity profile for the AQP1 overexpression (AQP1 OE) strain. In all image panels, dotted lines indicate the regions along which fluorescence intensity was quantified for the accompanying plots.
